# PHOTOTROPIN-mediated blue light signaling orients the asymmetry of *Marchantia polymorpha* spores

**DOI:** 10.64898/2026.03.22.713455

**Authors:** Johannes Roetzer, Radka Slovak, Eva-Sophie Wallner, Natalie Edelbacher, Beate Asper, Sebastian Deiber, Sebastian Seitner, Martin Colombini, Liam Dolan

## Abstract

Multicellular organisms produced by sexual reproduction develop from single cells and the asymmetry of these cells can define the orientation of the earliest developmental axes. The haploid multicellular stage of the plant, *Marchantia polymorpha,* develops from a single cell – the spore – that divides asymmetrically, producing an apical germ cell that generates the plant body and a smaller basal cell that differentiates as an anchoring germ rhizoid cell. We show that the orientation of this asymmetric cell division is controlled by an external, environmental cue – blue light – that is perceived by the photoreceptor PHOTOTROPIN and signals in an NCH1-dependent manner. This defines core elements of the mechanism by which a directional environmental signal orients cell division, which in turn orients the first axis of symmetry.

## Introduction

Land plants develop separate, multicellular haploid and diploid phases in their life cycle. The diploid phase develops from a zygote produced by the fertilization of a polarized egg (*1*). The polarity established in the egg is inherited by the zygote (*2*), which orients the first cell division and aligns the apical-basal axis of the embryo through a mechanism in which auxin modulates the activity of transcription factors in different regions of the embryo (*3*). The haploid phase develops from the cells produced by meiosis (spores). Spores are dispersed in an anhydrous state and become activated upon contact with water, and swell, forming spherical cells. Spore cells then divide asymmetrically, and this asymmetry in turn aligns the first body axis that defines the position of the first meristem from which the mature plant body develops (*4–6*). The cues that orient the spore cell asymmetry are unknown.

A dry *M. polymorpha* spore is approximately 7-10 µm in diameter and expands to about 20 µm within the first 24 hours after imbibition. After approximately 28 hours, the spore nucleus moves from the cell centroid to the cell cortex at the future basal pole. There, the nuclear membrane breaks down, the mitotic spindle forms, chromosomes are separated at mitosis, and a new cell wall forms near the basal pole during cytokinesis, producing a small cell near the basal pole and a larger cell which includes the entire apical hemisphere and most of the basal hemisphere. The large apical cell divides to form a population of cells from which a meristem forms, while the smaller basal cell differentiates into a germ rhizoid, which anchors the developing plant to the substrate (*7–9*). The movement of the nucleus to the basal pole requires reorganization of the microtubule and actin microfilament cytoskeletons, which provide both the direction and force for nuclear movement (*10–13*). However, neither the cue nor the molecular mechanism that orients spore cell asymmetry in *M. polymorpha* is known. Here, we report the discovery that a blue light cue perceived by the PHOTOTROPIN photoreceptor and its interaction partner NCH1 mediate the perception of this environmental signal.

## Results

### Light positively regulates the division of the *M. polymorpha* spore

Dry *M. polymorpha* spores (Figure 1A) become spherical with a diameter of approximately 7-10 µm, within 12 hours of imbibition (Figure 1D). By 24 hours after initiation of imbibition, spores were approximately 20 µm in diameter, with dense cytoplasm and a nucleus located near the centroid (Figure 1B, D). Spores divided asymmetrically after approximately 30 hours, forming a large apical germ cell and a smaller basal cell, separated by a new cell wall that was perpendicular to the apical-basal axis of the two-cell stage plant (Figure 1C-E) (*10, 14*). The larger daughter cell is generative and proliferates into a mass of morphologically similar cells that form the main body of a sporeling (Figure 1C), on which a meristem develops *de novo* 8-10 days after plating in white light (*15*). The basal cell differentiates as a tip-growing rhizoid cell.

**Figure 1.**
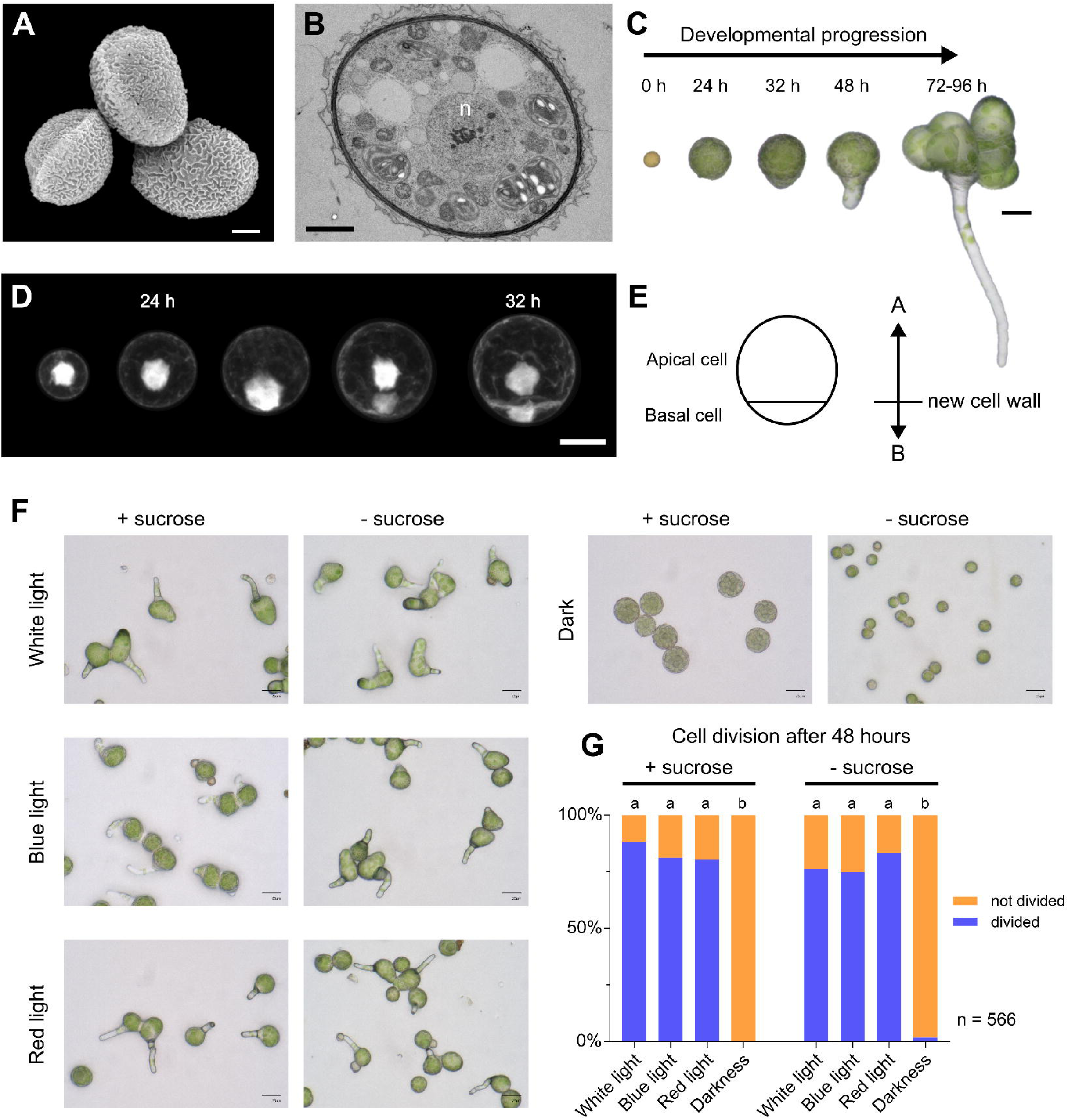
Light positively regulates the division of the *M. polymorpha* spore. **(A)** SEM image of dry *M. polymorpha* spores. Scale bar = 2 µm. **(B)** TEM image of a wild type spore 24 h after initiation of imbibition, showing dense cytoplasm and a nucleus (n) located near the cell centroid. Scale bar = 3 µm. **(C)** Representative bright-field images of spores at the indicated times after plating. **(D)** Visualization of nucleus position in spores expressing nuclear and plasma membrane markers (*pro*Mp*ROP:mScarletI-N7* and *pro*Mp*UBE2:mScarletI-*At*LTI6b*). Spores were imaged after 12, 24, 29, and 32 h. Maximum intensity projections (xy) of 3D z-stacks from five representative spores are shown. Scale bar = 10 µm. (**E)** Schematic of the two-cell spore stage following the first cell division, showing a larger apical cell and a smaller basal cell. The newly formed cell wall is oriented perpendicular to the apical-basal axis. **(F)** Spores grown for 2 days in darkness, white light, blue light, or red light on nutrient media with or without sucrose. Scale bars = 25 µm. **(G)** Quantification of spore cell division after 48 h. Proportions of dividing and non-dividing spores (total n = 566) are shown for each condition. Chi-square tests of independence revealed significant effects of light treatment both with sucrose (χ²(3) = 161.7, p < 0.0001) and without sucrose (χ²(3) = 72.1, p < 0.0001). Different letters indicate statistically significant differences (p < 0.05); identical letters indicate no significant difference.

To test the role of light in sporeling development, we grew spores in darkness, white light, red light, and blue light in nutrient media without a sucrose supplement. Spore cells did not divide if grown in darkness for up to 7 days (2-day data, Figure 1F-G, Figure S1A; 5-day data, Figure S1B; 7-day data, Figure S1C). By contrast, 76% of spore cells divided after two days in white light, 83% in red light, and 75% in blue light (Figure 1F-G, Figure S1A). These data indicate that light is required for cell division (*16*). To test if the lack of cell division in darkness was due to carbon starvation resulting from a lack of photosynthesis, we grew spores in nutrient media supplemented with 1% (w/v) sucrose (Figure 1F, Figure S1A). Dark-grown spores did not divide (0%) after two days, while spores grown in white (88%), red (80%), and blue (81%) light divided (Figure 1G). These data indicate that light is required to stimulate spore division in the first days after imbibition, and that monochromatic red or blue light alone is sufficient to initiate division. Dark-grown spores divided after five (73%) and seven days (86%) in darkness on sucrose-supplemented medium, indicating that spores can eventually divide in the absence of light (Figures S1B-C). From these data, we conclude that the first cell division of the spore is dependent on light at approximately 30 h.

### Blue light orients the first cell division

We reasoned that either light, gravity, or both would control the cell division orientation. To determine if light directs nuclear migration that positions and orients cell division, we developed a chamber in which spores were cultivated under unidirectional light. The larger apical cell developed on the side nearest the incident light, while the smaller basal cell developed on the relatively shaded side (opposite the incident light direction) (Figure 2A). The perpendicular orientation of the new cell wall relative to the incident light results in the apical cell being located on the illuminated side and the basal cell on the relatively shaded side. We measured the angles between the direction of incident light and the apical–basal axis (Figure S2A). The angles were close to 0°, indicating that the light vector aligns with the apical–basal axis in spores grown under unidirectional white light (Figure 2A). These data suggest that light orients the asymmetric cell division in the spore cell. However, in these experiments, the light and gravity vectors were aligned, making it impossible to distinguish between the two.

**Figure 2.**
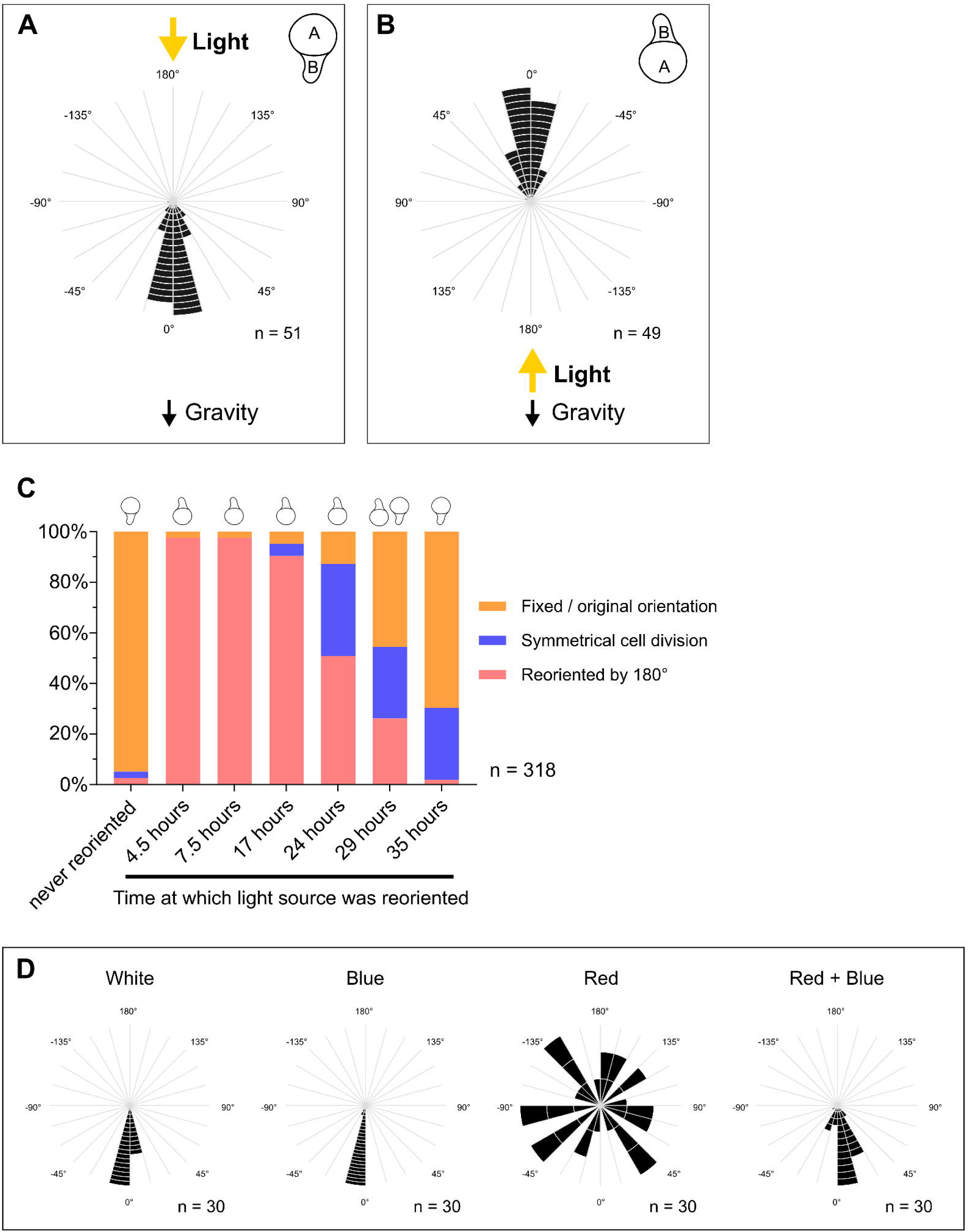
Blue light orients the first cell division. **(A)** Radial histogram of wild type spores (n = 51) grown for 48 h in unidirectional white light in 3D agar culture in chambered microscopy slides. Illumination from above aligned light and gravity vectors. The new cell wall formed perpendicular to the incident light, with the apical cell on the illuminated side and the basal cell on the relatively shaded side. **(B)** Radial histogram of wild type spores (n = 49) grown for 48 h in unidirectional white light with light and gravity vectors opposed (illumination from below). The new cell wall formed perpendicular to the incident light, and the basal cell developed on the relatively shaded side. These data indicate that light, not gravity, orients asymmetric division. **(C)** Reorientation experiments show that division orientation remains labile until shortly before division. Spores (n = 318) were grown in unidirectional white light, and the direction of light was reversed by 180° at different time points; all samples were analyzed after 48 h. In controls (never reoriented), the basal cell formed on the relatively shaded side. **(D)** Orientation assays under different light conditions show that blue light orients the first cell division. Spores were grown for 48 h in white, red, blue, or red+blue light in 3D agar culture. All orientation experiments were performed using wild type spores expressing the fluorescent marker *pro*Mp*SYP13A:mCitrine:*Mp*SYP13A* to visualize cell division.

To test if gravity contributes to the orientation of spore cell asymmetry (*17–19*), we aligned the light and gravity vectors at 180° to each other by shining light from below. If gravity defined the direction of asymmetry, we would expect the smaller basal cell to develop on the lower, illuminated side and the larger apical cell to develop on the upper, shaded side. On the other hand, if light oriented asymmetry, we would expect the smaller basal cell to develop at the upper shaded side and the large apical cell to develop at the lower illuminated side. When illuminated from below, the smaller basal cell formed at the upper shaded side, and the new cell wall was oriented perpendicular to the incident light (Figure 2B). These data do not support the hypothesis that gravity directs asymmetry. Instead, they indicate that light directs the development of cellular asymmetry that orients asymmetric division of the spore.

To understand when the light signal acts during the orientation of spore asymmetry, we tested the stability of light-induced asymmetry. We grew spores for varying durations in white incident light from one direction before changing the direction of the incident light by 180°. Measurements were taken after a total of 48 hours of light exposure for all treatments. In controls where light was applied from one direction for the duration of the experiment, cell division was asymmetric, and the smaller basal cell formed on the relatively shaded side. When the direction of light was reversed by 180° at different time points, up to 29 hours, the position of the smaller basal cell formed by asymmetric cell division was also reversed (Figure 2C, Figure S2B). This indicates that the light-directed asymmetry is labile and changes if the direction of incident light changes until approximately 29 hours. Reversal of the light direction at 24 and 29 hours resulted in a mixed population where the smaller basal cells developed at either pole, or in some cases, division was symmetrical, forming two equal-sized cells (Figure 2C, Figure S2B). Spore germination and cell division are asynchronous, and nuclear migration occurs between 24 and 29 hours (*10*). The mixed orientations observed at these late time points are consistent with developmental asynchrony within the spore population, such that individual spores were at different stages when the light direction was altered. Reversal of the light direction at 35 hours did not affect the position of the smaller basal cell – the smaller cell always developed on the shaded side relative to the direction of first illumination (Figure 2C, Figure S2B). Since asymmetric cell division occurs at around 30 hours, we conclude that the cell asymmetry remains labile until shortly before cell division.

To determine which light wavelengths direct cell wall orientation during the development of cellular asymmetry, we grew spores in white, red, and blue light and determined the position of the smaller basal cell and new cell wall orientation. For spore orientation assays, we built a custom illumination system (Figure S2C; Data S1, “Lightbox”) in which spores cultured in 0.4% agar in chambered microscopy slides were illuminated by directional light. This system allowed us to image and measure cell division orientation of spores in three-dimensional space. In white light, the smaller basal cell developed on the relatively shaded side, and the new cell wall was oriented perpendicular to the incident light (Figure 2D). In blue light, the smaller basal cell also developed on the relatively shaded side, and the new cell wall was oriented perpendicular to the incident light (Figure 2D). However, in red light, the smaller basal cell was randomly positioned, and the new cell wall was randomly oriented relative to the incident light (Figure 2D). When we grew spores in red light supplemented with blue light, they oriented similarly to those grown in monochromatic blue light, with the smaller basal cell forming on the shaded side and the new cell wall oriented perpendicular to the incident light, further underpinning the role of blue light in orienting spore cells (Figure 2D). We measured the angles between the direction of incident light and the apical-basal axis (Figure S2A) under the four tested light conditions: white, blue, red, and combined red and blue light. Significant directional clustering occurred in both white and blue conditions (Rayleigh test, p < 0.001, mean resultant length r = 0.99 and 0.98, respectively), as did the combined red and blue light treatment (p < 0.001, r = 0.94). In contrast, angles of the spores grown in red light were not clustered (p = 0.998, r = 0.009). Watson-Wheeler tests revealed significant differences in angular distributions among all groups (multi-group test p < 0.001), with all pairwise comparisons also showing statistically significant differences (p < 0.005).

To further test the role of light in orienting cell division, we determined the orientation of cell division in those spores that divided after 7 days in the dark on sucrose-supplemented media (Figures S1B-C). The position of the smaller basal cell was not aligned and was random as observed in spores grown for 48 hours in red light (Figure S2D). These data indicate that blue light provides the directional cue that orients the asymmetry of cell division. While red light stimulates cell division, it does not orient cell asymmetry.

### MpPHOT-mediated blue light sensing orients the first cell division

Because blue light was required to orient the asymmetric cell division in the spore (Figure 2), we hypothesized that a blue light receptor mediated this response (*20–22*). To identify the blue light photoreceptor responsible for orienting spore cell division, we generated transcriptomes from sporelings at 0, 15, and 30 hours after imbibition and performed RNA sequencing. mRNA levels of several blue light photoreceptors – *PHOTOTROPIN* (Mp*PHOT*), *CRYPTOCHROME* (Mp*CRY*), and *FLAVIN-BINDING KELCH REPEAT F-BOX PROTEIN* (Mp*FKF*) – were abundant (Figure 3A). We hypothesized that MpPHOTOTROPIN would regulate the direction of nuclear movement (*23*) because of its described role in directing chloroplast movement (*24–27*).

**Figure 3.**
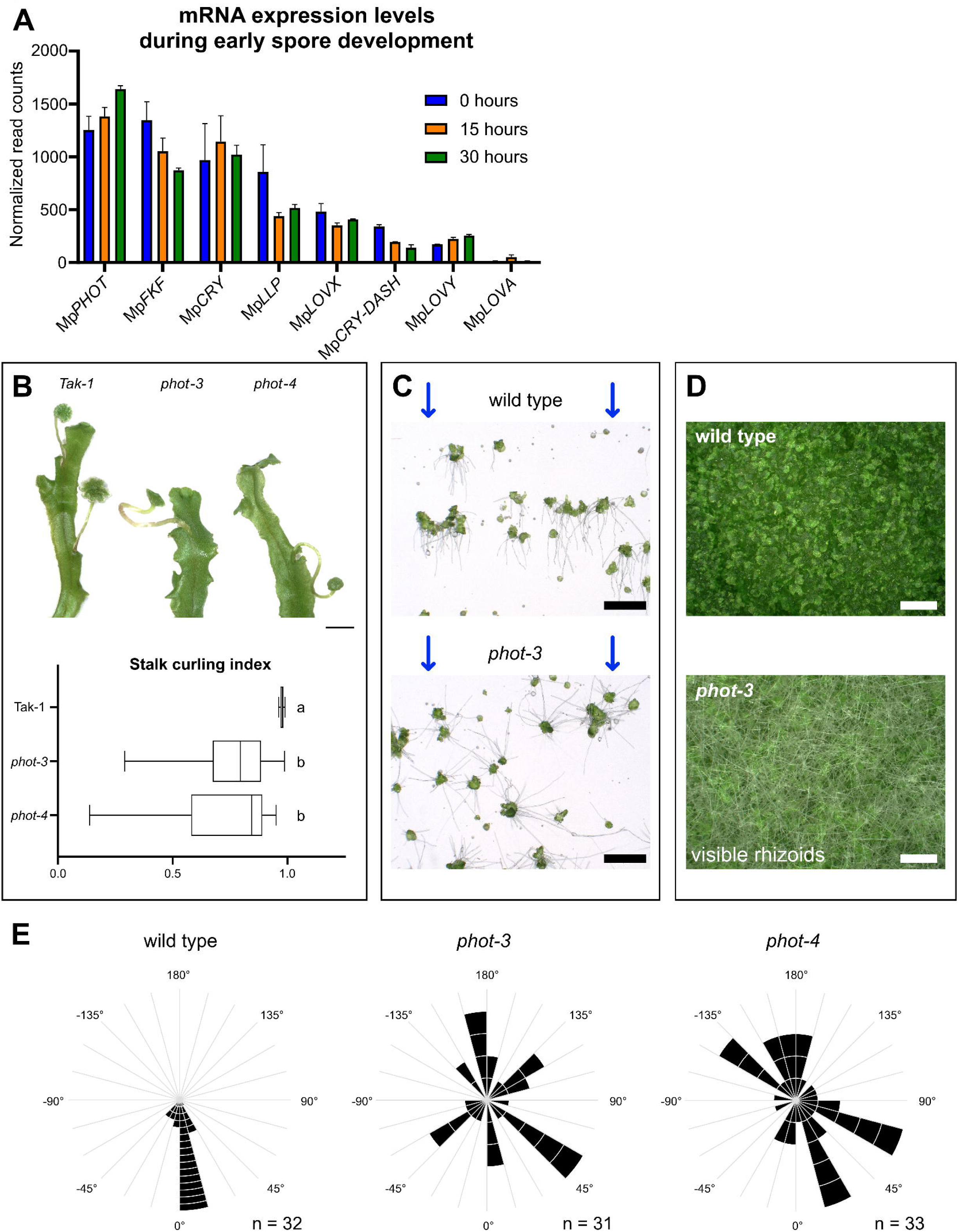
MpPHOT-mediated blue light sensing orients the first cell division. **(A)** Blue light receptor genes are expressed early during spore development. Shown are normalized mRNA read counts at 0 h (blue), 15 h (orange), and 30 h (green); two biological replicates per time point; mean with SD. **(B)** Positive phototropism of gametangiophore stalks is defective in Mp*phot* mutants. Stalk curling was quantified in 8-week-old plants grown under inductive conditions (n = 46 across three genotypes). Scale bar = 5 mm. Data were tested for normality (Shapiro-Wilk); as assumptions were not met, a Kruskal–Wallis test with Dunn’s multiple comparisons was performed. Different letters indicate significant differences (p < 0.05); identical letters indicate no significant difference. (**C)** Rhizoids of Mp*phot* mutant sporelings are defective in negative phototropism in blue light. In wild type, rhizoids of 10-day-old sporelings grew away from the blue light source, whereas mutant rhizoids grew in many directions. Scale bar = 1 mm. **(D)** Rhizoids of 11-day-old Mp*phot* mutant sporelings grew in many directions in white light. Under standard white-light conditions on horizontal plates, rhizoids were visible in mutants but not in wild type. Spores were densely plated and imaged from above. Scale bar = 2 mm. **(E)** MpPHOT is required to orient asymmetric cell division. Wild type and Mp*phot* mutant spores expressing *pro*Mp*SYP13A*:*mCitrine*:Mp*SYP13A* were grown for 48 h in continuous blue light. In wild type, the basal cell formed on the shaded side, and the new cell wall was oriented perpendicular to the incident light. In Mp*phot-3* and Mp*phot-4*, the position of the basal cell and cell wall orientation were random.

To test if Mp*phot* mutants are defective in blue light signaling, we characterized phototropic responses in independent mutant lines (*28*). Positive phototropism was defective in Mp*phot* mutants: wild type gametangiophore stalks grew straight and erect towards the light source. By contrast, Mp*phot* mutant gametangiophore stalks grew in different directions, resulting in curled stalk morphology (Figure 3B). Stalk curling was quantified using a stalk curling index defined as the ratio between the shortest and within-tissue distance from the stalk base to the base of the antheridial head. The index of straight stalks approaches 1, but is lower in curved stalks. The average index of wild type was ∼0.98, whereas the index of Mp*phot* mutants was ∼0.74 (Figure 3B). The defective positive phototropism in Mp*phot* mutants is consistent with the hypothesis that blue light-mediated phototropism is defective in these mutants.

Negative phototropism was also defective in Mp*phot* mutants. In wild type, rhizoids of 10-day-old sporelings grew away from the blue light source, while Mp*phot* mutant rhizoids were insensitive to light direction and grew in all directions (Figure 3C).

Similarly, under standard white-light conditions on horizontal plates, rhizoids were visible in 11-day-old Mp*phot* sporelings but not in wild type when densely plated spores were imaged from above (Figure 3D). The defective gametangiophore growth and defective rhizoid negative phototropism in Mp*phot* mutants demonstrate that MpPHOT is required for both positive and negative phototropic responses and that blue light signaling is defective in the Mp*phot* loss-of-function mutants.

To determine if MpPHOT is required to orient cell division asymmetry relative to the direction of incident blue light, we compared the position of the smaller basal cell and the orientation of the new cell wall that forms in unilateral light in wild type and Mp*phot* mutants. The smaller basal cell developed on the shaded side of the two-cell sporeling, and the new cell wall was oriented perpendicular to the direction of incident light in the wild type (Figure 2A, Figure 3E). By contrast, in Mp*phot* mutants, the smaller basal cell developed in random positions, and the orientation of the new cell wall was random with respect to the direction of light (Figure 3F). We measured the angles between the direction of incident light and the apical-basal axis. Wild-type clustered at consistently low angles, indicating that they all were oriented in the same direction (Rayleigh test, *p* < 0.001, mean resultant length *r* = 0.95), while Mp*phot-3* and Mp*phot-4* mutants were randomly oriented (*p* > 0.5, r = 0.12 and 0.14, respectively). Watson-Wheeler tests revealed significant differences in angular distributions between wild type and both mutants (*p* < 0.001), but not between Mp*phot-3* and Mp*phot-4* (*p* = 0.99). These data indicate that the position of the smaller basal cell and the orientation of the new cell wall were defective in Mp*phot* mutants. We conclude that MpPHOT is required for the development of blue light-directed cell asymmetry that orients asymmetric cell division of the spore.

### MpNCH1 mediates MpPHOT-dependent orientation of asymmetric cell division

To identify proteins involved in MpPHOT-mediated blue light signaling, we performed in vivo proximity labeling using a MpPHOT–miniTurbo(ID)-YFP fusion protein.

Biotinylated proteins from two independent *pro*Mp*PHOT*:Mp*PHOT*-*miniTurbo-YFP* lines were compared with those from a compartment control (*pro*Mp*PHOT*:Mp*PIP*-*miniTurbo-YFP*) and the wild-type control (Tak-1). All constructs were introduced into the same genetic background, Tak-1, by thallus transformation. Proteins biotinylated in the vicinity of MpPHOT were identified by mass spectrometry. This identified 547 to 550 proteins that were significantly enriched (log2FoldChange >2, p-value < 0.05) for two independently tested PHOT-miniTurbo-YFP lines against Tak-1 (Figure 4 A, Table S1). From the enriched candidates, we prioritized eleven proteins for further analysis based on their expression in the one-cell spore stage (≤30) transcriptomes (Figure 3A) and their predicted roles in signal transduction (candidates highlighted in Figure 4A). To test if any of the eleven proteins were required for blue light-oriented division of the spore cell, we generated a total of 137 mutant lines carrying loss-of-function mutations in the respective genes (Figure S4A).

**Figure 4.**
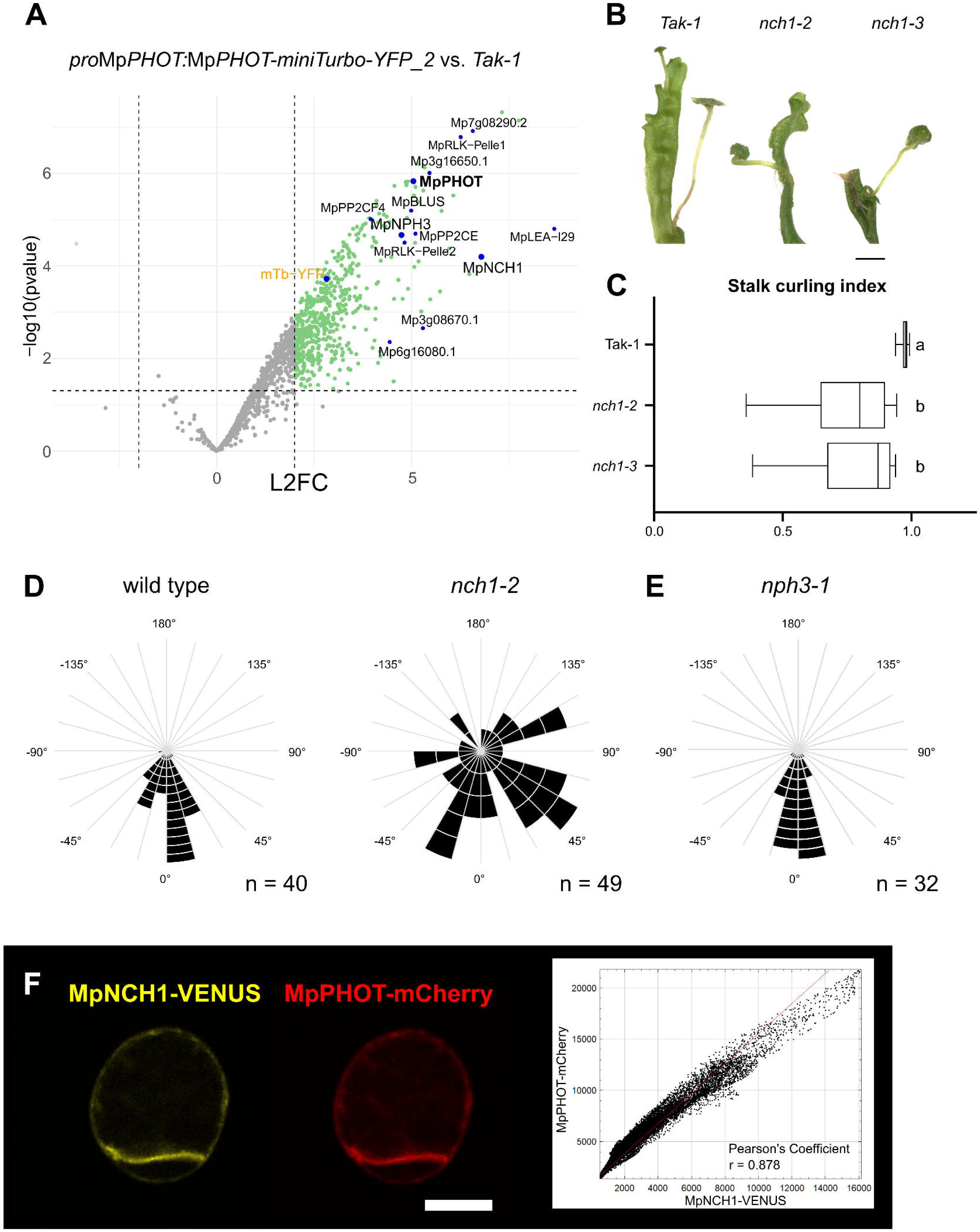
MpNCH1 mediates MpPHOT-dependent orientation of asymmetric cell division. **(A)** Proximity-labeling-based proteomics identifies proteins in close proximity to MpPHOT. MpPHOT–miniTurbo(ID)–YFP fusion proteins were used to biotinylate proteins in continuous white light, followed by mass spectrometry. The volcano plot shows differential enrichment relative to wild type (Tak-1) from triplicate samples (p < 0.05, logLJFC > 2). MpPHOT and miniTurbo–YFP are enriched as expected (self-labeling), together with MpNCH1, MpBLUS, MpNPH3, and additional proteins highlighted in the graph. Eleven proteins enriched in two independent lines and expressed during early spore development (0–30 h; spore transcriptomes, Figure 3) were selected for further analysis. Additional comparisons of independent experimental lines with compartment and wild-type controls are provided in Table S1. **(B)** Positive phototropism of gametangiophore stalks is defective in Mp*nch1* mutants. Scale bar = 5 mm. **(C)** Quantification of stalk curling in 8-week-old plants grown under inductive conditions (n = 37 across three genotypes). Data were tested for normality (Shapiro-Wilk); non-normal data were analyzed using a Kruskal-Wallis test with Dunn’s multiple comparisons. Different letters indicate significant differences (p < 0.05); identical letters indicate no significant difference. **(D)** MpNCH1 is required to orient asymmetric cell division. Wild-type and Mp*nch1* spores expressing *pro*Mp*SYP13A*:*mCitrine*:Mp*SYP13A* were grown for 48 h in continuous blue light. In wild type, the basal cell formed on the shaded side and the new cell wall was oriented perpendicular to the incident light, whereas both were random in Mp*nch1*. **(E)** MpNPH3 is not required for orientation of asymmetric cell division. Mp*nph3* spores grown under the same conditions showed wild type-like orientation. **(F)** MpNCH1–VENUS and MpPHOT–mCherry co-localize in spores. The cytofluorogram shows pixel intensity correlation from a single plane of an individual cell (thresholded; Gaussian blur, radius = 2). Signals are strongly correlated (Pearson’s r = 0.878), indicating overlapping localization. Spores were grown for 36 h on cellophane-covered medium, fixed, cleared, and imaged by confocal microscopy. Scale bar = 10 µm.

Next, we characterized light responses in the thallus and gametangiophores. While some developed defective thalli, only mutants in one gene – Mp*NRL PROTEIN FOR CHLOROPLAST MOVEMENT1* (Mp*NCH1*) – developed the curled gametangiophore phenotype characteristic of defective blue light signaling, as observed in the Mp*phot* mutants (Figure 4B). The stalk curling index of Mp*nch1* mutants was approximately 0.77 (Figure 4C) and similar to Mp*phot* mutants (Figure 3C), whereas the wild type developed an index close to 1 (∼0.97; Figure 4C). These data indicate that blue light signaling is defective in Mp*nch1* mutants.

We then compared the position of the smaller basal cell and new cell wall orientation in Mp*nch1* mutants and wild type. In wild type, the new basal cell formed on the shaded side, and the new cell wall was oriented perpendicular to the incident light (Figure 4D). However, in Mp*nch1* mutants, the smaller basal cell developed in random positions relative to the direction of illumination, and the new cell wall was randomly oriented (Figure 4D). We measured the angles between the direction of incident light and the apical-basal axis. The orientation angles of wild type are clustered (Rayleigh test, *p* < 0.001, *r* > 0.9), whereas the orientation of the axis is largely random with only weak residual directionality (*r* = 0.34) in Mp*nch1* mutants.

Consistent with this, the angular distributions of wild type and Mp*nch1* differ significantly (Watson–Wheeler test, *p* < 0.001). These data indicate that Mp*NCH1* is required for blue light-directed asymmetry that orients asymmetric cell division in the spore.

MpNCH1 is a member of a family of NRL (NPH3/RPT2-like) proteins that have been implicated in MpPHOT-mediated blue light signaling (*29*). MpNONPHOTOTROPIC HYPOCOTYL3 (MpNPH3) is also an NRL protein family member that functions in blue light signaling (*30*). To test if MpNPH3 is required for blue light-oriented spore cell division, we generated lines carrying mutations in Mp*NPH3* and compared the positioning of the smaller basal cell and new cell wall orientation in Mp*nph3* loss-of-function mutants with wild type. Both the position of the small basal cell and the orientation of the new cell wall in the Mp*nph3* spores were wild type-like (Figure 4E). The orientation angles are clustered in Mp*nph3*, similar to wild type (Rayleigh test, *p* < 0.001, *r* > 0.9). These data indicate that Mp*NPH3* is not required for the orientation of cell asymmetry in the *M. polymorpha* spore.

If MpPHOT and MpNCH1 interact during the establishment of cell asymmetry in the spore, we predicted that the proteins should co-localize during this process. To test this hypothesis, we generated marker lines expressing *pro*Mp*NCH1:*Mp*NCH1-VENUS* and *pro*Mp*PHOT:*Mp*PHOT-mCherry* in the same spore. Spores were grown for 24 h and 36 h under standard white light, then fixed and cleared to remove chloroplast autofluorescence. MpPHOT-mCherry and MpNCH1-VENUS co-localized at the cell surface (Figure 4F, Figure S4B). These data are consistent with MpPHOT and MpNCH1 acting in the same pathway to regulate blue light-oriented cell asymmetry in the *M. polymorpha* spore.

## Discussion

We discovered that MpPHOT-mediated blue light signaling aligns the cellular asymmetry that orients the first division of the spore and sets up the first developmental axis of the *M. polymorpha* body plan. MpPHOT-mediated blue light signaling that directs the orientation of cellular asymmetry requires MpNCH1 but not the related MpNPH3 NRL protein (*29*). These data demonstrate that the orientation of cell asymmetry and subsequent body axis orientation depend on an external blue light cue from the environment. Such externally directed cellular asymmetry positions the germ rhizoid on the shaded side of the spore. This rhizoid cell anchors the plant to the substrate during its early development (*4*). The apical cell is on the illuminated side, divides and forms a photosynthetically active sporeling that develops a meristem from which the plant body develops (*15*).

In land plants, the diploid zygote develops within the maternal tissues in which the egg cell formed: within the archegonium in bryophytes, lycophytes, monilophytes, and gymnosperms, and within the embryo sac in angiosperms (*1*). Signals from the parental tissues polarize the egg and zygote (*3, 31*). By contrast, haploid spores that form in bryophytes and monilophytes develop in the environment and independent of the plant on which they are produced. The cues that orient development do not come from surrounding tissues (*11*). In free-sporing plants, including several fern species, blue light has been shown to inhibit spore germination, as reviewed in Suo et al (*32*). Instead, we show here that blue light orients the symmetry-breaking event that occurs in spores in the first 30 hours of growth before cell division. This cell asymmetry orients the first cell division, the first body axis, and subsequently the negative phototropic growth of the germ rhizoid. In this way, MpPHOT-mediated blue light signaling orients the development of the *M. polymorpha* body, tethering the plant to its substrate.

The demonstration that both MpPHOT and MpNCH1 are required to direct cell asymmetry in the spore cell suggests that a canonical MpPHOT signaling pathway regulates this process. In addition to MpNCH1, proximity labeling revealed several proteins previously linked to phototropin signaling, including a BLUS-like protein (*33*) and multiple kinases and phosphatases (Figure 4A, Fig S4A). These associations are consistent with engagement of a broader phosphorylation-dependent signaling framework. Nevertheless, our functional analyses identify MpNCH1 as a key downstream component required for orienting spore cell asymmetry. This discovery reveals how plants exploit a directional environmental cue to align activities in the single totipotent cell from which the multicellular haploid plant develops.

## Acknowledgements

We thank Katharina Jandrasits and Magdalena Mosiolek for general laboratory support. We are grateful to the ProTech Facility of the Vienna BioCenter for cloning the MpNCH1 CRISPR constructs, and to the Vienna BioCenter Core Facilities for continuous support. This includes the BioOptics Facility, with special thanks to Thomas Lendl for assistance with co-localization image analysis, as well as the Plant Sciences Facility, Mechanical Engineering Center, Electron Microscopy Facility, Proteomics Facility, Molecular Biology Service, Media Lab, Lab Support, and the administrative staff at the VBC.

Transmission electron microscopy (TEM) of spores was performed with Marlene Brandstetter at the Electron Microscopy Facility, Vienna BioCenter Core Facilities (VBCF), member of the Vienna BioCenter (VBC), Vienna, Austria. Scanning electron microscopy (SEM) was carried out with Daniela Gruber at the Core Facility Cell Imaging and Ultrastructure Research (CIUS), University of Vienna, a member of Vienna Life-Science Instruments (VLSI). RNA sequencing was performed at the Wellcome Trust Centre for Human Genetics, University of Oxford. Proteomic analyses were carried out by the Proteomics Facility at IMP/IMBA/GMI using the Vienna BioCenter Core Facilities (VBCF) instrument pool.

We thank Haonan Bao, Zohar Meir, and Victoria Spencer for critically reading the manuscript, and Zohar Meir for assistance with uploading the transcriptome data to the repository.

## Author contributions

Conceptualization: JR, RS, LD

Methodology: JR, RS, ESW, SS, MC, LD

Investigation: JR, RS, ESW, NE, BA, SD

Visualization: JR, ESW

Funding acquisition: LD

Project administration: LD

Supervision: JR, LD

Writing – original draft: JR, LD

Writing – review & editing: JR, ESW, LD

## Data, code, and materials availability

Requests for resources and further information should be directed to the lead contact, Liam Dolan.

All data reported in this paper will be made available by the lead contact upon request following publication of the peer-reviewed paper. The raw and processed data for the spore transcriptomes will be available in the NCBI Gene Expression Omnibus (GEO) repository upon publication of the peer-reviewed paper. The mass spectrometry proteomics data have been deposited in the ProteomeXchange Consortium via the PRIDE partner repository (*34, 35*) with the dataset identifier PXD072764 and will be available upon publication of the peer-reviewed paper.

Requests for plasmids and transgenic lines generated in this study should be directed to the lead contact. Recipients are required to provide appropriate documentation, such as import permits, for the transfer of transgenic material.

## Materials and Methods

### Plant material and growth conditions

Wild-type accessions of *Marchantia polymorpha*, Takaragaike-1 (Tak-1, male) and Takaragaike-2 (Tak-2, female) (*36*), together with the respective mutant lines, were cultivated on sterile solid medium. The growth medium consisted of half-strength Gamborg’s B5 (*37*) supplemented with 0.5 g/L MES hydrate, 1% (w/v) sucrose, and 1% (w/v) plant agar, adjusted to pH 5.5 before autoclaving. The plants were cultivated at 23 °C under continuous white light illumination in standard LED growth chambers at the institute or in growth chambers manufactured by Poly Klima Climatic Grow Systems (Hofmann Kühlung Company; firmware version 03.10.10(22), controller: WAGO 750-8212 PFC200 G2 2ETH R). For dark treatment controls, plates were wrapped twice with aluminium foil, transferred into a light-impermeable container, and maintained in the same growth chamber.

Unless otherwise stated in the respective figures, the following light regimes were applied: standard continuous white light at a photon flux density (PFD) of 50-60 μmol m⁻² s⁻¹; blue light at approximately 30-40 μmol m⁻ ² s⁻¹ (λmax = 450 nm); and red light at approximately 80-90 μmol m⁻ ² s⁻ ¹ (λmax = 660 nm). Dark controls were prepared as described above and kept at 23 °C in the growth chamber. Light intensities were quantified using a Spectral PAR Meter PG100N (UPRtek).

For gametangiophore induction, plants were transferred to soil in SacOLJ Microbox containers containing an autoclaved 3:1 mixture of Neuhaus N3 compost and vermiculite, and grown under 50 μmol m⁻ ² s⁻¹ white light supplemented with 45 μmol m⁻² s⁻¹ far-red light in a long-day regime (16 h light / 8 h dark) at 23 °C.

### Scanning electron microscopy of dry *Marchantia polymorpha* spores

Sporangia from a cross between Tak-1 and Tak-2 were collected and dried over silica gel at room temperature for 3-4 weeks. Dry *M. polymorpha* spores were mounted on SEM stubs and sputter-coated with gold for 30 seconds using a JEOL JFC-2300HR sputter coater to generate a conductive surface layer.

Samples were imaged using a JEOL IT300 scanning electron microscope equipped with a LaB6 filament. Secondary electron and backscattered electron detectors were used for image acquisition. SEM imaging was performed at the Core Facility Cell Imaging and Ultrastructure Research (CIUS), University of Vienna, member of the Vienna Life-Science Instruments (VLSI).

### Transmission electron microscopy of *Marchantia polymorpha* spores

*Marchantia polymorpha* spores were cultured on cellophane-covered medium plates under continuous white light conditions for 24 hours after plating. The spores were then washed off the cellophane and immediately pre-fixed in 0.5% (w/v) paraformaldehyde and 0.05% (v/v) glutaraldehyde in 0.1 M PHEM buffer for 10 minutes. Samples were washed twice in PHEM buffer, transferred to Gamborg medium supplemented with 10% (w/v) BSA (fraction V), and cryo-immobilized using a Wohlwend HPF Compact 01 high-pressure freezer.

Freeze-substitution was performed in a Leica EM AFS automated system over 3 days at -90 °C in acetone containing 0.1% (w/v) tannic acid and 0.5% (v/v) glutaraldehyde. After substitution, samples were warmed to -20 °C over 12 h, washed in pre-cooled acetone, and incubated in 2% (w/v) osmium tetroxide in acetone for 24 h for infiltration and post-fixation. Samples were then warmed at ∼5 °C/h to 4 °C, rinsed, incubated for 2 h at 4 °C, brought to room temperature, and washed further in acetone.

Samples were infiltrated with Agar 100 epoxy resin through a graded resin:acetone series and polymerized in pure resin at 60 °C for 48 hours. Ultrathin sections (70 nm nominal thickness) were cut using a Leica UCT ultramicrotome and post-stained with 2% (w/v) uranyl acetate and Reynolds’ lead citrate.

TEM imaging was performed on an FEI Morgagni 268D transmission electron microscope equipped with a tungsten filament emitter and a MegaView III CCD camera (Olympus-SIS). The instrument was operated at 80 kV (nominal maximum 100 kV). Images were acquired using Morgagni software (v3.0) and iTEM (v5.0). Digital images were recorded at 7100x magnification with an image size of 1376×1032 pixels (16-bit), corresponding to a pixel size of 9.151 nm.

TEM imaging of *Marchantia* spores was carried out at the Electron Microscopy Facility, Vienna BioCenter Core Facilities (VBCF), member of the Vienna BioCenter (VBC), Vienna, Austria.

### Bright-field imaging of plants and spores

Plants and spores were imaged using a Keyence VHX-7000 digital bright-field microscope equipped with VH-Z00R/Z00T and VH-ZST lenses and a VHX-7020 camera. Where necessary, non-specific background was removed, and images were cropped to appropriate dimensions for figure preparation.

### Spore generation, culture, and imaging

Crosses between male and female plants were performed by collecting sperm from antheridiophores in sterile water for 15 min and transferring the suspension onto archegoniophores. Fully developed sporangia were collected 4-5 weeks after fertilization, placed in sealed polypropylene microboxes (OV80+OVD80) with micropore tape, and dried over silica gel (*36, 38*). After a 4-week desiccation period, sporangia were stored at -70 °C until use.

For germination, frozen sporangia were thawed and gently crushed in 0.1% (w/v) sodium dichloroisocyanurate (NaDCC; Sigma-Aldrich, Cat. No. 218928) for 1-2 min. Spores were pelleted (13,000 x g, 3 min), washed twice with sterile water, and resuspended in sterile water.

Spores were plated on sterile medium consisting of half-strength Gamborg B5 supplemented with 0.5 g/L MES hydrate (pH 5.5), containing 1% sucrose or no sucrose, and solidified with 1% plant agar or 1% phytoagar. Cultures were grown at 23 °C under continuous illumination. For dark controls, plates were wrapped twice in aluminium foil, placed in a light-tight box, and maintained in the same growth chamber.

For spore orientation assays, wild type and mutant spores expressing *pro*Mp*SYP13A:mCitrine:*Mp*SYP13A* were embedded in half-strength B5 medium (1.5 g/L B5, 0.5 g/L MES, 1% sucrose, pH 5.5) solidified in 0.4% plant agar. The agar medium was cooled to near-solidification before mixing with spores to generate a homogeneous 3D culture, which was solidified in chambered microscopy slides (Ibidi, Cat. No. 80286). Slides were placed in a custom-built illumination system (“lightbox”).

All procedures involving spores for light assays were performed under dim green light in a sterile laminar flow hood.

### Custom-built illumination system (Lightbox) for spore orientation experiments

Spore orientation assays were performed using the custom-built Lightbox system described in this section. This setup enables controlled directional illumination of spores in 0.4% 3D agar culture in chambered microscopy slides.

#### Software and control

The Lightbox control software was developed mainly by Sebastian Seitner (PlantS Facility, GMI Vienna). The system comprises four independent chambers, each equipped with a dedicated sample holder and LED module (Figure S2C, Data S1). All modules can be controlled independently.

Illumination regimes are defined as schedules and uploaded to the device via USB-C in JSON (JavaScript Object Notation) format. Each schedule specifies the sequence, duration, and intensity of light conditions for each chamber. The uploaded schedules are selected via an integrated user interface consisting of a rotary control and LCD display.

Each module contains 16 high-power LEDs (OSRAM), covering four spectral ranges: white (3500 K), blue (455 nm), red (657 nm), and far-red (727 nm).

Light intensity is regulated using pulse width modulation (PWM) at ∼30 kHz. PWM enables stable low-intensity output without shifting emission spectra. LEDs of each color are arranged in two electrically independent groups. These groups are driven with a 180° phase offset. At intensities above 50%, at least one group remains active at any time. This reduces temporal fluctuations in light delivery.

#### Hardware

The Lightbox hardware was developed by Martin Colombini and the Mechanical Engineering Center (Vienna BioCenter).

The system consists of a base plate, a control unit, and a light-tight enclosure containing four independent chambers. Each chamber fits one two-well chambered coverslip from IBIDI (Cat. No. 80286).

The enclosure includes a rear panel supporting the LED boards, a front panel with an access door, structural side panels, internal partitions, and top covers. Samples are inserted through the front door and positioned precisely within each chamber. Illumination parameters are set via the external control unit.

Thermal control is facilitated through active airflow: a top-mounted fan draws air from the base and exhausts it upwards, cooling the LED assemblies. The rear LED compartment is covered by a removable panel to maintain airflow while allowing access for maintenance. Each chamber is additionally ventilated by side-mounted fans. Air enters through a filtered inlet, passes through a light-blocking labyrinth, and exits through a corresponding outlet. These structures prevent external light contamination.

Optical isolation between chambers is ensured by internal partitions, diaphragm elements, and sealing interfaces between the front panel and door. These features prevent light leakage and cross-illumination.

The structural components (base, front, rear, and side panels) are constructed from rigid PVC. Covers, partitions, and the door are made from anodized aluminium. Internal rails are aluminium with a sandblasted finish to reduce reflections. Interior panels consist of black acrylic. Components are assembled primarily using stainless steel screws.

#### Previous setup for spore orientation experiments

Initial (re)orientation and gravity experiments were performed using a simplified setup. Unidirectional illumination was provided by single LEDs mounted on a breadboard. To expose spores to unidirectional light, chambered microscopy slides (Ibidi, Cat. No. 80286) containing spores were placed inside a 3D-printed cover box, in which all sides except one were enclosed by a light-proof black cover. This configuration allowed light to enter from only one side of the chamber, creating unidirectional illumination.

The entire setup was placed in an opaque enclosure (a black cardboard box) for the duration of the experiment. Light intensity was measured using a Spectral PAR Meter PG100N (UPRtek).

### Spore orientation quantification – angle calculation

Angles between the light vector and the apical-basal axis were measured using FIJI and visualized in RStudio. Two-dimensional or z-stacked confocal images were acquired from wild-type and mutant spores expressing the fluorescent marker *pro*Mp*SYP13A:mCitrine:*Mp*SYP13A* to visualize cell division. Images were oriented such that the direction of incident light corresponded to the negative y-axis.

In FIJI, the transverse cell wall of asymmetrically dividing sporelings was identified manually. Using the multi-point tool, the two junctions where the transverse wall meets the spore outline were marked sequentially, and XY coordinates were exported for each image. Angles were calculated in RStudio using the exported coordinate files. The apical-basal axis was defined as the vector orthogonal to the transverse cell wall. The angle between this axis and the light vector (negative y-axis) was then computed.

All calculated angles from individual measurements for each genotype and experiment were used to generate binned radial histograms. Visualization was performed using the R packages circular, plotrix, and ggplot2.

### Fixation, clearing, and cell wall staining of spores

Spores expressing nuclear and plasma membrane markers (*10*) (*pro*Mp*ROP:mScarletI-N7* and *pro*Mp*UBE2:mScarletI-*At*LTI6b*; Figure 2D) and spores expressing fluorescent reporters for co-localization (*pro*Mp*NCH1:*Mp*NCH1-VENUS* and *pro*Mp*PHOT:*Mp*PHOT-mCherry*; Figure 4F) were cultivated on cellophane-covered standard medium with sucrose for the indicated time periods. Cellophane with spores was removed and submerged in 10% formalin containing 0.1% Brij L23 for 30 min. Fixed sporelings were pelleted (7,000 x g, 3 min), washed in 1x PBS, and cleared in ClearSee α solution (10% w/v xylitol, 15% w/v sodium deoxycholate, 25% w/v urea, supplemented with 6.3 mg/mL sodium sulfite anhydrous) for approximately one week in the dark with gentle agitation.

Cleared sporelings were mounted in ClearSee α and imaged by confocal microscopy (Zeiss LSM 880) using a 40x/1.2 LD LCI Plan-Apochromat water/glycerol DIC objective with silicone immersion oil matched to the refractive index of ClearSee α. For co-localization analysis (Figure 4F), single-channel z-stacks (VENUS and mCherry) were processed in ImageJ/Fiji (*39*), and co-localization was quantified using JACoP (*40*).

For cell division analysis under different light conditions (Figure 1G; Figures S1A-C), the same fixation and clearing protocol was used with the addition of 0.2% Renaissance SR2200 applied overnight prior to imaging. The dye for cell wall staining was excited at 405 nm, and emission was collected between 420-500 nm using an upright widefield microscope. Images were acquired with an sCMOS camera, and cell divisions were manually scored in FIJI (*39*).

### RNA-seq and differential expression analysis of early spore development

To assess gene expression during early spore development, spores were harvested in two biological replicates at three timepoints: 0, 15, and 30 hours after plating on standard medium plates covered with cellophane. For each sample, six wild type sporangia (derived from a cross of Tak-1 and Tak-2) were collected, and surface-sterilized with 1% NaDCC, followed by washing three times with sterile water. The spore suspension was plated and maintained under continuous white light (PFD 50-60 µmol m⁻ ² s⁻ ¹) for the indicated durations.

For collection, the cellophane with adhered spores was peeled from the medium, flash-frozen in liquid nitrogen, and ground to a fine powder using a mixer mill (RETSCH MM 400) at 30 Hz for 4 minutes. Total RNA was extracted using the RNeasy Plant Mini Kit (Qiagen) according to the manufacturer’s instructions, followed by Turbo DNase treatment to remove residual genomic DNA.

RNA sequencing was performed at the Wellcome Trust Centre for Human Genetics, University of Oxford, using an Illumina HiSeq 4000 platform generating 75 bp paired- end reads. Each sample was sequenced in five technical replicates, with replicates distributed across five independent sequencing lanes. Raw reads were returned as fastq.gz files. Reads passing quality control (Phred +33 quality score >30 assessed with FastQC) were trimmed by removing the first and last 10 bases, and adapters were removed using Trimmomatic (v0.32) (*41*). Paired-end reads were interleaved. Residual bacterial rRNA sequences were filtered with SortMeRNA (v2.1b) (*42*).

Reads were error-corrected using BayesHammer within SPAdes (v3.10.1) (*43*) and aligned to the *Marchantia polymorpha* reference transcriptome (v3.1) (*44*) using Salmon (*45*).

Differential gene expression analysis was performed with DESeq2 (*46*). Transcript-level quantifications were summarized to gene-level counts using the tximport algorithm, and technical replicates were collapsed using the collapseReplicates function in DESeq2 package (*46*) in R. Gene annotations were obtained from the *Marchantia polymorpha* reference genome (v3.1) (functional annotation downloaded from https://marchantia.info/download/v31/). Pairwise comparisons were conducted, with an adjusted p-value (padj) <0.05 considered significant. Genes with log2FoldChange <0 were classified as downregulated, >0 as upregulated, and NA indicated no significant change.

Between 0 and 15 hours, 2,953 genes were upregulated, 2,947 downregulated, and 13,317 unchanged. Of the 2,953 upregulated genes, 360 were further upregulated, 108 downregulated, and 2,485 unchanged at 30 hours. Of the 2,947 downregulated genes, 128 were upregulated, 97 further downregulated, and 2,722 unchanged at 30 hours. Among the 13,317 unchanged genes between 0 and 15 hours, 290 were upregulated, 85 downregulated, and 12,942 remained unchanged at 30 hours.

### Mutant generation using CRISPR/Cas9 mutagenesis

CRISPR/Cas9 mutagenesis was performed as described in detail in Roetzer et al. (*28*), including sgRNA design and target site selection, plasmid generation, bacterial transformation, generation of transgenic lines, *Agrobacterium*-mediated transformation of *Marchantia polymorpha* sporelings, and mutant screening and genotyping.

In brief, CRISPR/Cas9-mediated mutagenesis was used to generate mutant lines for Mp*NCH1* (Mp5g07060) and Mp*NPH3* (Mp5g17270). Single guide RNAs (sgRNAs) were designed using CHOPCHOP (*47*) or manually for regions not listed among the top hits, based on gene sequences from marchantia.info (MpTak v6.1r2) (*48*). Each sgRNA consisted of a 20-nt target sequence followed by a canonical NGG PAM, and candidate sgRNAs were checked for uniqueness against the *M. polymorpha* genome.

sgRNA duplexes were cloned into OP-076 vectors (L2_lacZgRNA-Cas9-CsA) from the OpenPlant kit (*49*), carrying dual antibiotic resistance markers. Recombinant plasmids were introduced into *E. coli* DH5α cells, verified by Sanger sequencing, and purified for transformation. Mutant lines were generated using spores derived from a cross between Tak-1 and Tak-2 and *Agrobacterium*-mediated transformation with strain GV3101 (*36*). Transformants were selected on appropriate antibiotic-containing media.

Genomic DNA from transformants was extracted and PCR-amplified across gRNA target sites. The PCR products were sequenced using Sanger sequencing to identify mutations, and lines harboring mutations predicted to generate premature stop codons were selected. Mutant lines were propagated, and genotypes were repeatedly confirmed.

### Stalk curling index

Stalk curling was quantified by calculating a stalk curling index (SCI). For this analysis, individual stalks bearing an antheridial head were imaged under standardized conditions. For each genotype, four independent plants were used.

To determine the SCI, two distances between the stalk base and the base of the antheridial head were measured. First, the ‘real length’ was determined by tracing the within-tissue path along the curvature of the stalk from its base to the base of the antheridial head. Second, the ‘shortest distance’ was measured as the linear distance between these same two reference points by drawing a straight line.

The stalk curling index was calculated as the ratio of ‘shortest distance’ to ‘real length’. An SCI value approaching 1 indicates a straight stalk, whereas lower values reflect increasing degrees of curvature. Length measurements were performed using the measurement software of the Keyence microscope VHX-7000.

### Sporeling orientation assay

Spores were surface-sterilized as described above and resuspended in sterile water (one sporangium per 500 µL). For vertical blue-light assays (Figure 3C, Figure S3A), 25 µL of the suspension was evenly distributed onto square plates containing standard growth medium (½-strength B5 Gamborg with 0.5 g/L MES hydrate, 1% (w/v) sucrose, 1% (w/v) plant agar, pH 5.5). All handling prior to illumination was performed under dim green light. After spreading spores with a sterile cell spreader, plates were air-dried for 15 min in a laminar flow hood.

To restrict directional illumination, plates were wrapped twice in black, light-absorptive aluminium foil, leaving only the upper edge of the plate exposed. Plates were positioned vertically in a climate chamber (Poly Klima Climatic Grow Systems;

Hofmann Kühlung; firmware 03.10.10(22), controller WAGO 750-8212 PFC200 G2 2ETH R) at 23 °C under continuous blue light (∼30–40 µmol m⁻² s⁻¹; λmax = 450 nm) for 10 days.

For white-light sporeling orientation assays (Figure 3D), spores were sterilized and handled as described above, except that 100 µL of suspension was plated per square plate. Plates were incubated horizontally under continuous white light at 23 °C. Images were acquired after 11 days using a Keyence microscope VHX-7000 as detailed above.

### miniTurbo-based proximity labeling (*50*) and mass spectrometry

Transgenic Tak-1 *Marchantia polymorpha* lines expressing MpPHOT-miniTurbo-YFP (*pro*Mp*PHOT*:Mp*PHOT*-*miniTurbo*-*YFP*_line-1 and line-2) and a plasma membrane control (*pro*Mp*PHOT*:Mp*PIP-miniTurbo*-*YFP*), as well as the Tak-1 wild-type background, were grown from gemmae on half-strength B5 Gamborg medium supplemented with 1% sucrose, 1% plant agar, and 0.5 g/L MES hydrate (pH 5.5). Plants were grown for 10 days under standard white light on plates covered with cellophane to facilitate rapid thallus harvesting.

#### Proximity labeling under standard continuous white light

Ten-day-old thalli were incubated in 100 μM biotin dissolved in liquid half-strength B5 Gamborg medium for 16 h. To stop the biotinylation, plant material was strained and rinsed three times with 200 ml ice cold water, patted dry with paper towels and flash-frozen in liquid nitrogen. The frozen plant material was ground into a fine powder by a mortar and aliquoted into technical triplicates of 500 μL plant powder per sample. Proteins were extracted by adding 1 ml of extraction buffer (50 mM Tris pH 7.5, 150 mM NaCl, 0.1% SDS, 1% NP-40, 0.5% Na-deoxycholate, 1 mM EDTA, 1 mM EGTA, 1 mM DTT, 20 µM MG-132, 1x cOmplete, 1 mM PMSF), incubating for 15 min on a rotor wheel at 4° C. To digest cell walls, 1 µl Lysonase (Millipore) was added per sample, followed by another 15 min incubation at 4° C before sonication in a water bath at high settings for 4x 30 sec with 90 sec breaks. After sonication, samples were centrifuged for 15 min at 4° C, 15000 x g and the supernatants were desalted using PD-10 columns (GE Healthcare) to remove free biotin. The eluate was collected in 5 ml LoBind tubes (Eppendorf) and mixed with 200 μl Dynabeads MyOne Streptavidin C1 (Invitrogen) slurry that was activated according to the manual and pre-washed with extraction buffer. Streptavidin beads were incubated with the extracts for 16 h at 4 °C on a rotating wheel. Dynabeads were separated on a magnetic rack in 1.5 ml LoBind tubes (Eppendorf) and washed 4x with 1.5 ml cold equilibration buffer, followed by washes with 1x with cold wash buffer I (equilibration buffer with 250 mM NaCl + 1% SDS), 1x with cold equilibration buffer, 2x with cold wash buffer II (equilibration buffer with 250 mM NaCl), 2x with cold wash buffer III (equilibration buffer with 500 mM NaCl), 1x with cold equilibration buffer and 2x with Urea wash buffer (2 M Urea, 50 mM Tris pH 7.5, room temperature) as also described in Mair et al (*51*) and Wallner et al (*52*). Beads were resuspended in 1 ml cold 50 mM Tris pH 7.5 and transferred into a new 1.5 ml LoBind tube and all buffer was removed using a magnetic rack. Beads were frozen at -70°C for further processing.

#### Proximity labeling under dark and blue-light conditions

The same MpPHOT transgenic lines were grown and adapted in complete darkness for 24 h. Plants were divided into three conditions: 1) labeling in darkness, 2) blue-light activation immediately upon transfer, and 3) continuous blue-light exposure for 4 h before labeling. Biotin treatment was performed with 200 μM dissolved in liquid half-strength B5 Gamborg medium for 4 h. Further processing of plant material was done as described for white light conditions.

#### On-bead digestion

Beads were resuspended in 60 μL 100 mM ammonium bicarbonate with 600 ng Lys-C (Fujifilm Wako) and incubated for 4 h at 37 °C on a thermoshaker (1200 rpm). The supernatant was reduced with 10 mM TCEP for 30 min at 60 °C and alkylated with 40 mM MMTS for 30 min at room temperature. Digestion was completed with 600 ng trypsin (Trypsin Gold, Promega) overnight at 37 °C. Digests were acidified with 10 μL 10% TFA and submitted to LC-MS/MS.

#### nanoLC-MS/MS analysis

Peptides were separated on a Vanquish Neo UHPLC system coupled to an Orbitrap Exploris 480 mass spectrometer with FAIMS pro interface and Nanospray Flex source. Trap and analytical columns were PepMap Acclaim C18 (5 μm, 100 Å) with dimensions 5 mm × 300 μm ID (trap) and 500 mm × 75 μm ID (analytical). Peptides were eluted at 230 nL/min using a 120-min linear gradient from 2% to 35% acetonitrile in 0.1% formic acid. Data were acquired in data-dependent acquisition mode (DDA) with full scans at 60,000 resolution, m/z 350–1200, and MS/MS scans of the most abundant ions with HCD at 30% collision energy. Compensation voltages of -45, -60, and -75 V were applied with cycle times of 0.9 s (CV -45 and -60) and 0.7 s (CV -75). Precursor ions of charge 2-6 were selected for fragmentation, with dynamic exclusion of 45 s.

#### Data processing and analysis

RAW files were processed in Proteome Discoverer 2.5 with MSAmanda 2.0. Peptides were searched against the UniProt *Marchantia polymorpha* database (34,956 sequences) and common contaminants. Fixed modification: beta-methylthiolation of cysteine; variable modifications: methionine oxidation, phosphorylation (S/T/Y), deamidation (N/Q), N-terminal pyro-glutamate, and N-terminal acetylation. FDR was set to 1% at PSM and protein levels using Percolator, and a minimum Amanda score of 150 was applied. Proteins were quantified with IMP-apQuant (MaxLFQ algorithm) and normalized using iBAQ with sum normalization across samples. Match-between-runs was applied, and proteins were filtered to require at least three quantified peptides per protein.

#### Volcano plot generation

Differential enrichment analysis was performed comparing each transgenic line to wild type and the control line. Volcano plots were generated in RStudio using packages ggplot2 and ggrepel. Significantly enriched proteins were determined by a p-value < 0.05 and a log2FoldChange > 2.

#### Proximity labelling - data availability

Mass spectrometry proteomics data from the proximity labelling experiments have been deposited to the ProteomeXchange Consortium via the PRIDE partner repository with dataset identifier PXD072764 and will be available upon publication of the peer-reviewed paper.

### Quantification and statistical analysis

Sequence assemblies and plasmid maps were generated using CLC Genomics Workbench v9.5.1 (Qiagen). Gene sequences were obtained from MarpolBase (https://marchantia.info/) (*48*). All experiments were repeated with 3-5 independent biological replicates. Sample sizes are indicated in the respective figure legends.

Data analysis and visualization were carried out using RStudio (v2022.07.2, Build 576) and GraphPad Prism (v8.1.1). Statistical analyses were performed in RStudio and GraphPad Prism.

**Figure S1. Light positively regulates spore division in *M. polymorpha*. Related to Figure 1F-G**.

**(A)** Overview images of spores grown for 2 days in darkness, white, blue, or red light on nutrient media with or without sucrose. Each overview image shows several spores used for quantification. Cell division occurred under all light conditions but not in darkness. Scale bars = 50 µm. **(B, C)** Quantification of spore cell division under white light and darkness after 5 days (B; n = 148) and 7 days (C; n = 123), with and without sucrose. For both time points, light treatment had a significant effect on cell division with sucrose (5 days: χ²(1) = 4.56, p = 0.033; 7 days: χ²(1) = 3.98, p = 0.046; chi-square test) and without sucrose (5 and 7 days: Fisher’s exact test, p < 0.0001). Different letters indicate statistically significant differences (p < 0.05); identical letters indicate no significant difference.

**Figure S2. Quantification and experimental setup for spore division orientation assays. Related to Figure 2**.

**(A)** Schematic of cell division orientation measurement. When the new cell wall (black) is oriented perpendicular to the light vector (yellow), the apical–basal axis (green) aligns with the light vector (left). When the cell wall is not perpendicular to the light vector, the axes are misaligned (right). The angle (blue) between the light vector and the apical–basal axis was measured for all spores used in orientation assays. **(B)** Radial histograms of spores used for reorientation experiments (Figure 2C). Only asymmetrically dividing spores were included. Light direction was reversed at 4.5 h (n = 40), 7.5 h (n = 40), 17 h (n = 40), 24 h (n = 35), 29 h (n = 41), and 35 h (n = 40), with a non-reoriented control (n = 38). All spores were imaged and quantified after 48 h total illumination. **(C)** Image of the custom-built illumination system (“lightbox”) used for spore orientation assays. The open-front view shows four units for chambered slides enabling unidirectional illumination. Details are described in *Materials and Methods*. **(D)** Radial histogram of spores dividing after 7 days in darkness on sucrose-supplemented medium (n = 35). Cell wall orientation was random.

**Figure S3. Rhizoids of Mp*phot-4* mutant sporelings are defective in negative phototropism in blue light. Related to Figure 3C**.

**(A)** Rhizoids of wild-type sporelings grew away from the blue light source, while Mp*phot-4* mutant rhizoids grew in multiple directions. Sporelings were grown for 10 days in continuous blue light on vertical plates with illumination from above. Scale bar = 1 mm.

**Figure S4. Selection of candidate genes and co-localization of MpNCH1–VENUS and MpPHOT–mCherry in the undivided spore. Related to Figure 4**.

(A) A total of 137 loss-of-function mutant lines were generated for selected genes using CRISPR–Cas9.

(B) MpNCH1–VENUS and MpPHOT–mCherry co-localize in spores before the first cell division. Representative single-plane confocal images of an undivided spore are shown, including individual channels and the merged image. Co-localization analysis revealed a strong correlation between the signals (Pearson’s r = 0.909), indicating overlapping subcellular localization. Spores were grown for 24 h on cellophane-covered medium, then fixed, cleared, and imaged by confocal microscopy. Scale bar = 10 µm.

## Supplementary Material

Figures: S1 to S4

Table S1: MS_table

Data S1: Lightbox

