## Supplementary figures and images for "PHOTOTROPIN-mediated blue light signaling orients the asymmetry of *Marchantia polymorpha* spores"

### Fig S1

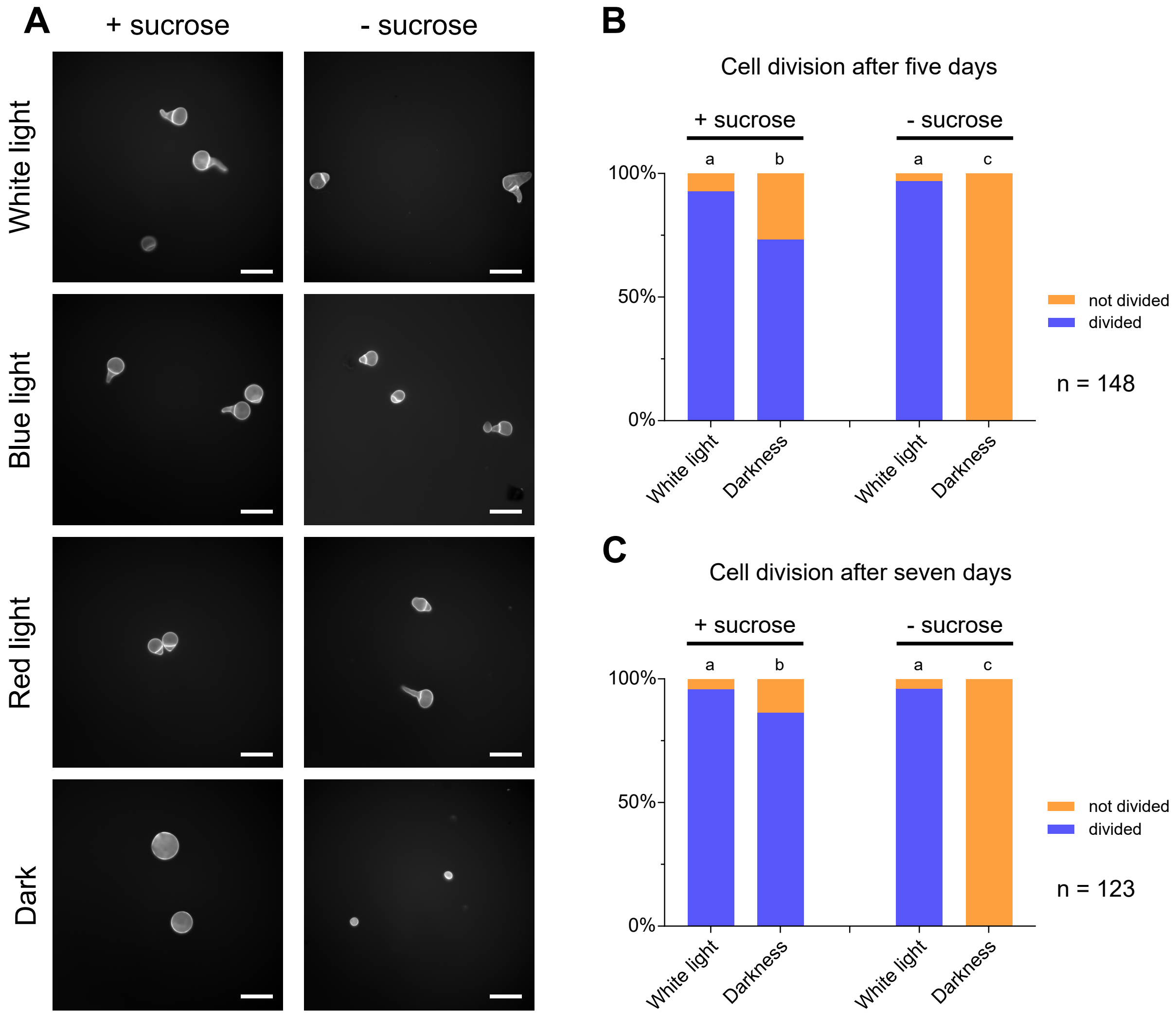

### Fig S2

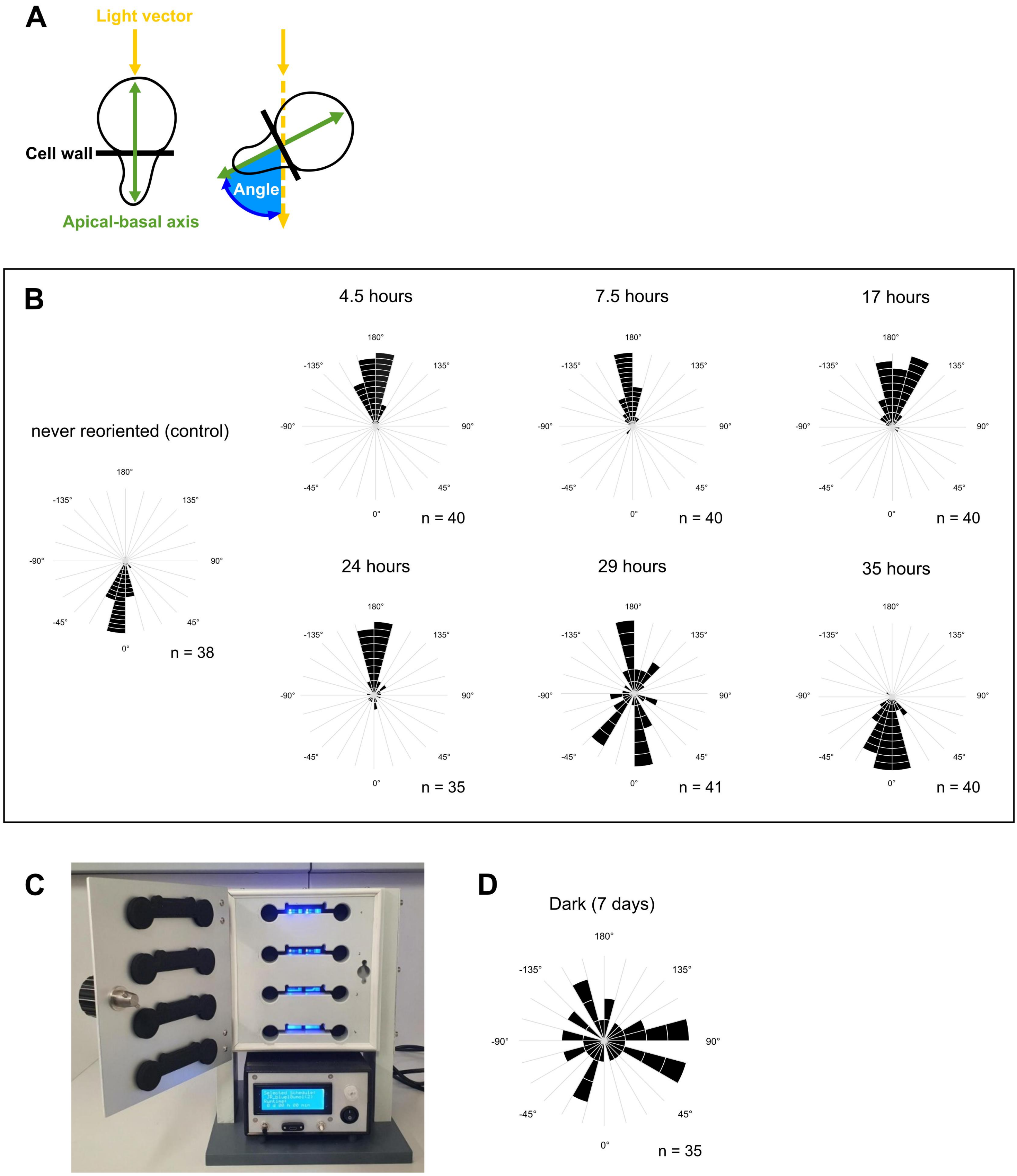

### Fig S3

#### Figure S3

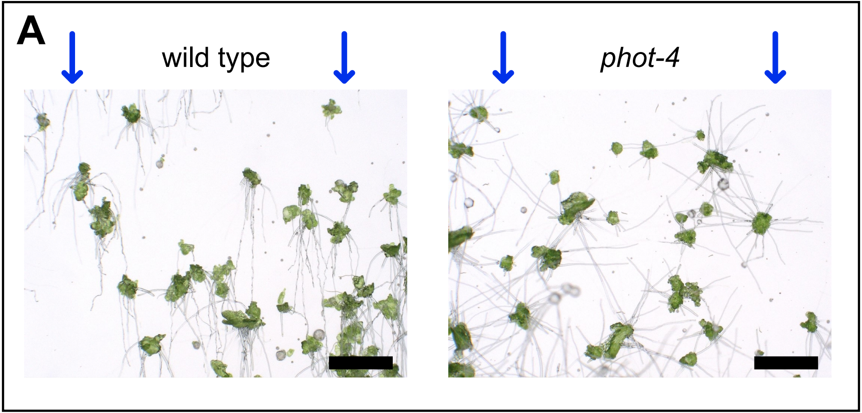
