## Supplementary material for "PHOTOTROPIN-mediated blue light signaling orients the asymmetry of *Marchantia polymorpha* spores": Fig S4

### Figure S4

## A

| Identifier | Available data / predicted function | No. of mutant lines |
|---|---|---|
| Mp5g07060 | Mp*NCH1* | 11 |
| Mp8g10870 | *BLUS*-like protein | 17 |
| Mp4g17810 | *LEA-like29* | 5 |
| Mp6g16080 | Kinesin light chain, TPR domain | 40 |
| Mp3g08670 | Serine/threonine kinase | 5 |
| Mp3g18020 | Receptor-like kinase | 3 |
| Mp3g05350 | Mp*RLK-Pelle_RLCK-II* receptor-like kinase | 3 |
| Mp3g16650 | *CDC2*-related protein kinase | 20 |
| Mp4g20380 | Mp*PP2C_E,* phosphatase | 7 |
| Mp5g14150 | Mp*PP2C_F4*, phosphatase | 1 |
| Mp7g08290 | *unknown* | 25 |

## B

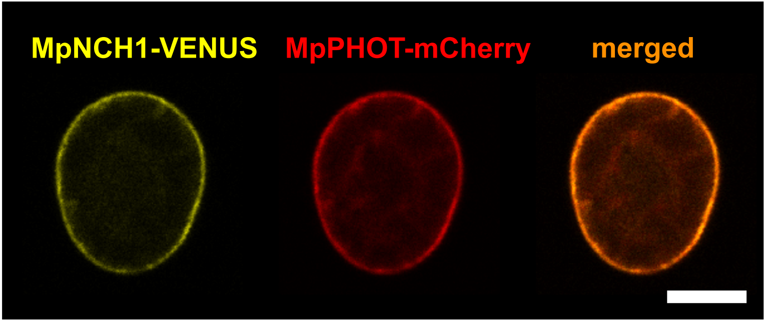
